# TrkC has distinct spatiotemporal dynamics compared to TrkA and TrkB

**DOI:** 10.1101/2025.11.13.688325

**Authors:** Ryan T. Duffy, Stephen J. Hill, Chloe J. Peach

## Abstract

Neurotrophins (NGF, BDNF, NT-3, NT-4) are critical regulators of neuronal development and have been implicated in a range of neurodegenerative disorders. They signal through three tropomyosin receptor kinase receptors (TrkA, TrkB and TrkC). TrkC can exist as an alternately spliced isoform (TrkC_KID_) that includes an exon-16 derived kinase insert domain (KID). Here, we have utilised FRET and BRET biosensors to compare the spatiotemporal dynamics of TrkA, TrkB and TrkC. TrkA and TrkB displayed robust concentration-dependent ERK signalling, dimerisation and trafficking, and also provided evidence for receptor activation by non-canonical ligands. While the KID-lacking TrkC_ΔKID_ showed ERK signalling comparable to TrkA and TrkB, TrkC_KID_ displayed markedly reduced downstream ERK signalling. Interestingly, TrkC_ΔKID_ and TrkC_KID_ exhibited slower trafficking away from the cell surface than TrkA or TrkB, but a strong ability to dimerise. Overall, these findings reveal that TrkC possesses unique regulatory mechanisms that may provide opportunities for tailored neurotrophic drug discovery.

## Introduction

Neurotrophins are a highly conserved family of growth factors critical for the development and maintenance of the nervous system, including nerve growth factor^1^ (NGF), brain-derived neurotrophic factor^2^ (BDNF), neurotrophin-3^3^ (NT-3), and neurotrophin-4/5^4^ (NT-4). Neurotrophins signal through the tropomyosin receptor kinase (Trk) family of receptor tyrosine kinases (RTKs), comprising TrkA, TrkB and TrkC, for which the canonical ligands are NGF/TrkA^5^, BDNF/TrkB, NT-4/TrkB^6^ and NT-3/TrkC^7^. Trk receptors are notably homogenous and have emerged as important therapeutic targets for neurodegenerative disorders, with TrkB particularly implicated across various disorders of the central nervous system (CNS)^8,9^. However, TrkC also plays a critical role in the CNS, with TrkC knockout leading to profound deficits in neurons *in vivo*, leading to death in the early postnatal period^10^. TrkC has also been shown to co-express with TrkB^11^, suggesting that while efforts to date have focused on modulating TrkB signalling, TrkC may also play an important role that remains less well characterised in these systems.

Growth factor stimulation promotes receptor dimerisation, leading to the auto- and trans-phosphorylation of key intracellular tyrosine residues^12^. This enables the recruitment of key intracellular adaptor proteins, initiating downstream signalling cascades such as mitogen-activated protein kinase (MAPK/ERK), phosphoinositide 3-kinase (PI3K/AKT) and phospholipase Cγ signalling pathways^12^. TrkC, however, can also function as a ‘dependence receptor’ that promotes apoptosis in the absence of ligand^13^ through caspase-mediated TrkC cleavage at aspartic acid residues D495 and D641^14,15^. This can be prevented by the binding of canonical ligand NT-3, which instead drives kinase signalling^14,15^. Together, these signalling events coordinate a range of neuronal processes including survival, synaptic plasticity and circuit refinement^9,16^. Following ligand binding, TrkA and TrkB are known to undergo receptor endocytosis in a clathrin-dependent manner^17^, facilitating retrograde transport in endosomes along axons to defined subcellular compartments^18^, from which TrkA^19^ and TrkB^20^ can continue to signal. While the trafficking and intracellular signalling properties of TrkA and TrkB have been well studied, the biological role of TrkC remains less well characterised, particularly in systems that lack co-expression of other Trk receptors.

Neurotrophic receptors can exist in multiple isoforms, such as the truncated variants of TrkB and TrkC that lack the intracellular kinase domain – TrkB.T1^21^ and TrkC.T1^22^. There are additional isoforms of TrkC generated through alternative splicing of exon 16, encoding a kinase insert domain (KID)^23,24^. This 14 amino acid (aa) insert is positioned immediately adjacent to the major autophosphorylation site (Y705, Y709, and Y710) in the activation loop^24^.

The 839aa TrkC_KID_ is expressed in a tissue- and developmental-stage specific manner, where similarly to its insert-lacking 825aa counterpart (TrkC_ΔKID_), both are detected in the same neuronal populations. Functionally, TrkC_KID_ has been shown to have impaired signalling compared to TrkC_ΔKID_^23,24^, however the extent to which this 14aa sequence affects receptor dynamics remains unknown.

Despite our understanding of the role of Trk receptors in physiology^9,16^, there are limited direct comparisons between the spatial and temporal pharmacology of TrkC with TrkA and TrkB expressed in isolation. To monitor Trk receptor pharmacology in an isolated system, resonance energy transfer-based approaches can be used to monitor real-time signalling such as fluorescence resonance energy transfer (FRET) and bioluminescence resonance energy transfer (BRET)^25^. These techniques rely on the transfer of energy from a donor molecule to an acceptor molecule when within proximity (<10 nm), allowing the quantitative analysis of protein-protein interactions with temporal resolution at 37°C. Building on previous studies with other RTKs using these methods^26^, including at TrkA^27^, we utilised FRET- and BRET-based approaches to systematically characterise and compare the real-time pharmacology of TrkA, TrkB, TrkC_KID_, and TrkC_ΔKID_ . By directly comparing downstream ERK signalling, dimerisation and trafficking in response to NGF, BDNF, NT-3 and NT-4, we identified both shared and divergent features of Trk receptor dynamics, with TrkC being distinct with respect to its ability to undergo ligand-induced dimerisation and endocytosis, irrespective of its ability to signal.

## Methods

### Cell Culture and materials

HEK293T cells were cultured using Dulbecco’s Modified Eagle Medium (DMEM) containing 10% foetal bovine serum (FBS) (Sigma Aldrich, US) and 1% penicillin streptomycin antibiotic (Sigma Aldrich, US). HEK293T cells were incubated at 37°C in a humidified incubator with 5% CO_2_. Cells were monitored daily and were split using a ratio of 1:10 or 1:15 to maintain a suitable growth confluency. Once cells reached 90% confluency, HEK293T cells were seeded at a density of 25,000 cells per well in a standard 96-well white clear bottom plate (#655089, Greiner Bio-One) for BRET experiments. For FRET, a standard 96-well black clear bottom plate (#G655209, Greiner Bio-One) was used. Hanks’ Balanced Salt Solution (HBSS), with calcium and glucose was purchased from Thermo Fisher Scientific (#14025092). HEPES buffer (H23830), Bovine Serum Albumin (A9418), and Phorbol 12,13-dilbutyrate (PDBu) (P1269) were all purchased from Sigma Aldrich. Furimazine was bought from Promega Corporation (N1110). SNAP-Surface Alexa Fluor 488 was bought from (New England Biolabs, S9129S). All neurotrophic growth factors were bought from Alomone Labs, including NGF (#N-245), BDNF (#B-250), NT-3 (#N-260), and NT-4 (#N-270). Coelenterazine Purple was bought from NanoLight technology (#369-1). Polyethyleneimine (PEI) was purchased from Polysciences (24765).

### Generating DNA Constructs

N-terminal NanoLuc-tagged and SnapTag-tagged TrkA, (NM_001012331), TrkB (NM_006180), TrkC (TrkC-TrkC_KID_; NM_002530, UniProt Q16288-1) and TrkC (TrkC_ΔKID_; NM_002530 with deletion of residues 712–725, UniProt Q16288-3) were generated in the pFN28K expression vector. Each construct included the IL-6 signal peptide fused to the N-terminus of either nanoluciferase (NanoLuc) or SnapTag respectively, followed by a GSSGAIA linker and the coding sequence of the respective Trk receptor. C-terminal *Renilla* luciferase (Rluc8)-tagged TrkA, TrkB, TrkC_ΔKID_, and TrkC_KID_ were cloned in a vector encoding the full length Trk receptor, followed by a HA-tag (YPYDVPDYA), and then the Rluc8 sequence. In addition, cleavage site mutations D495N and D641N were introduced into TrkC KID for all tagging formats using site-directed mutagenesis with a Q5 High-Fidelity DNA Polymerase (#E0554S, NEB).

### DNA Transfections

For all studies, following 24h, HEK293T cells were serum starved prior to transfection using DMEM without serum supplementation to remove growth factors in FBS. Cells were then transfected with transfection reagent polyethyleneimine (PEI) at ratio of 1:6 DNA to PEI. Cells were incubated for 48 hours to allow cells to express the transfected constructs prior to assay.

To quantify trafficking, HEK293T cells were transfected with each Trk receptor tagged on the C-terminus with Rluc8 either TrkA-HA-Rluc8, TrkB-HA-Rluc8, TrkC_ΔKID_-HA-Rluc8, TrkC_KID_-HA-Rluc8 (WT), TrkC_KID_-HA-Rluc8 (D495N), or TrkC_KID_-HA-Rluc8 (D641N) (20ng/well), as well as a marker of the plasma membrane tagged with Renilla green fluorescent protein (RGFP), RGFP-CAAX (20 ng/well). To quantify dimerisation, HEK293T cells were transfected with DNA encoding Trk receptors tagged on the N-terminus with a NanoLuc (NanoLuc-Trk, 10 ng/well) and a SnapTag (SnapTag-Trk, 25 ng/well). To quantify ERK signalling, HEK293T cells were transfected with nuclear EKAR CFP YFP (40 ng/well; Addgene, #18681) and N-terminal SnapTag-Trk (20 ng/well).

### Nuclear ERK FRET Assays

All assays used HBSS with calcium and glucose (#14025092 Thermo Fisher Scientific, US) supplemented with HEPES buffer (pH 7.4) with 0.1% Bovine Serum Albumin (BSA). On the day of the assay 48h post-transfection, cells were incubated in assay buffer and left to incubate for at least 30 minutes before stimulating with increasing agonist concentrations of NGF, BDNF, NT-3 or NT-4 (1 pM-30 nM). Phorbol 12,13-dibutyrate (PDBu, 10 µM) was used as a positive control and the buffer solution was used as a vehicle treatment. FRET reads were measured in the PHERAstar microplate reader (BMG Labtech) using a “FI 430 530 480” optic module. Donor excitation was provided at 430 nm (CFP), donor emission was recorded at 480 nm, and acceptor emission (YFP) was recorded at 530 nm following donor excitation. Following 5 baseline reads, increasing concentrations (1 pM-30 nM) of NGF, BDNF, NT-3 or NT-4 were added. The plate was then read every 63 seconds for a further 20 minutes. FRET ratio was calculated as ratio (530nm/480nm) of the two emission signals.

### Dimerisation BRET Assay

HEK293T cells were transfected with DNA encoding Trk receptors tagged at the N-terminus with either a luminescent donor, NanoLuc, or a SnapTag. Following 48h serum starvation and transfection, cells were incubated with SNAP-Surface Alexa Fluor 488 to label each receptor at the plasma membrane. Cells were pre-incubated with SnapTag (0.25 µM, 1 hour) at 37°C. Cells were then washed in assay buffer to remove unbound substrate. Cells were incubated with furimazine (NanoLuc substrate; 600X dilution) for 5 minutes at 37°C. BRET was measured using the PHERAstar FSX microplate reader (BMG Labtech). Firstly, 5 pre-ligand BRET reads were completed, allowing a “baseline BRET” to be calculated to indicate the proximity of tags prior to ligand stimulation. Next NGF, BDNF, NT-3 or NT-4 were added at increasing concentrations (1 pM-30 nM). The plate was then read every 45 seconds for a further 20 minutes using a BRET1 plus filter (donor wavelength of 475 nm and acceptor wavelength 535 nm). BRET ratios were calculated by dividing the emission intensity of the acceptor wavelength by the donor wavelength emission intensity, thus providing a ratiometric change in BRET.

### Trafficking BRET Assay

Following 48h serum starvation and transfection, assay buffer was added to each well, followed by Coelenterazine Purple (BRET substrate for Rluc8; 400X dilution). Each plate was incubated at 37°C for 5 minutes to allow the enzymatic reaction to occur. BRET reads were measured in the PHERAstar FSX microplate reader (BMG Labtech) using a BRET2 filter, (donor wavelength of 370–450 nm and acceptor wavelength 500–540 nm). 5 pre-ligand BRET reads were completed, allowing “baseline BRET” to be calculated. Following 5 baseline reads, increasing concentrations (1 pM-30 nM) of NGF, BDNF, NT-3 or NT-4 were added. The plate was then read every 74 seconds for a further 25 minutes. BRET ratios were calculated as explained in the dimerisation assay.

### Data Analysis

Data were analysed using GraphPad Prism (10.1.2). Data were baseline-corrected to the 5 reads prior to ligand addition, then baseline-corrected to changes in vehicle over time. For ERK signalling, data were then normalised to the maximal response of the positive control (PDBu, 100%), expressed as a percentage relative to vehicle (0%). Concentration-response data were fit using non-linear regression analysis as log vs. response (three parameters) “Y=Bottom + (Top-Bottom)/(1+10^(LogEC50-X)” to determine EC_50_ values. Concentration–response curves from the Trk trafficking assay were inverted to represent receptor endocytosis driven by growth factor agonism. To compare kinetic values, data from experiments using 30 nM NGF, BDNF, NT-3, and NT-4 were analysed by non-linear regression using a one-phase association model: “Y = Y_0_ + (Plateau – Y_0_) (1 – e^-Kx^)” to determine Kobs. Data points prior to 0 minutes were excluded, as no ligand had been added at that time. For ERK signalling specifically, data between 2 and 15 minutes were used to ensure an appropriate kinetic fit that captured a single association phase. Significance was determined on concentration-response data using ordinary two-way ANOVA followed by Dunnett’s multiple comparisons test, with all experimental groups compared to vehicle (α = 0.05). Significance was accepted at *p*<0.05 (*), *p*<0.01 (**), *p*<0.001 (***), and *p*<0.0001 (****). pEC50 and K_obs_ data was firstly assessed for normality using a Shapiro-Wilk test. From this, significance from normal data was determined using an ordinary one-way ANOVA followed by Dunnett’s comparisons test. Non-normal data was assessed using a Kruskal-Wallis test followed by Dunn’s multiple comparisons test or Welch’s T-test depending on if the analysis was conducted between 3 or more groups or just between 2 groups. Analysis was performed comparing non-canonical ligand vs canonical pairing for each receptor (e.g. TrkA/NGF vs TrkA/BDNF, TrkA/NT-3, and TrkA/NT-4), with significance *p*<0.05 (*), *p*<0.01 (**), *p*<0.001 (***), and *p*<0.0001 (****). Analysis was also performed across NT-3, compared to TrkC_ΔKID_/NT-3 (i.e TrkC_ΔKID_/NT-3 vs TrkA/NT-3, TrkB/NT-3, TrkC_KID_/NT-3) with significance denoted as *p*<0.05 (^†^), *p*<0.01 (^††^), *p*<0.001 (^†††^), and *p*<0.0001 (^††††^).

## Results

### Trk signalling in response to canonical and non-canonical neurotrophic factors

To determine the downstream consequences of ligand stimulation, we used a FRET-based assay to assess the level of ERK phosphorylation following ligand stimulation. Specifically, an EKAR sensor localised to the nucleus was used due to the proximity of the sensor to transcriptional events. This sensor enabled live-cell detection of ERK activity within this defined subcellular compartment (*Figure 1A*). When ERK is activated, it phosphorylates the substrate domain on EKAR; this induces a conformational change that brings the donor and acceptor fluorophores into close proximity, inducing an increase in FRET^28^. Phorbol 12,13-dibutyrate (PDBu) was used as a positive control as an activator of Protein Kinase C, thus leading to ERK activation that is independent from Trk activity. ERK activity was first assessed in HEK293T cells transfected with EKAR only to observe any baseline signalling in the absence of Trk receptor transfection. When stimulated with 30 nM of NGF, BDNF, NT-3 or NT-4, this led to no detectable increase in FRET signal compared to vehicle throughout (*Figure 1B*).

**Figure 1.**
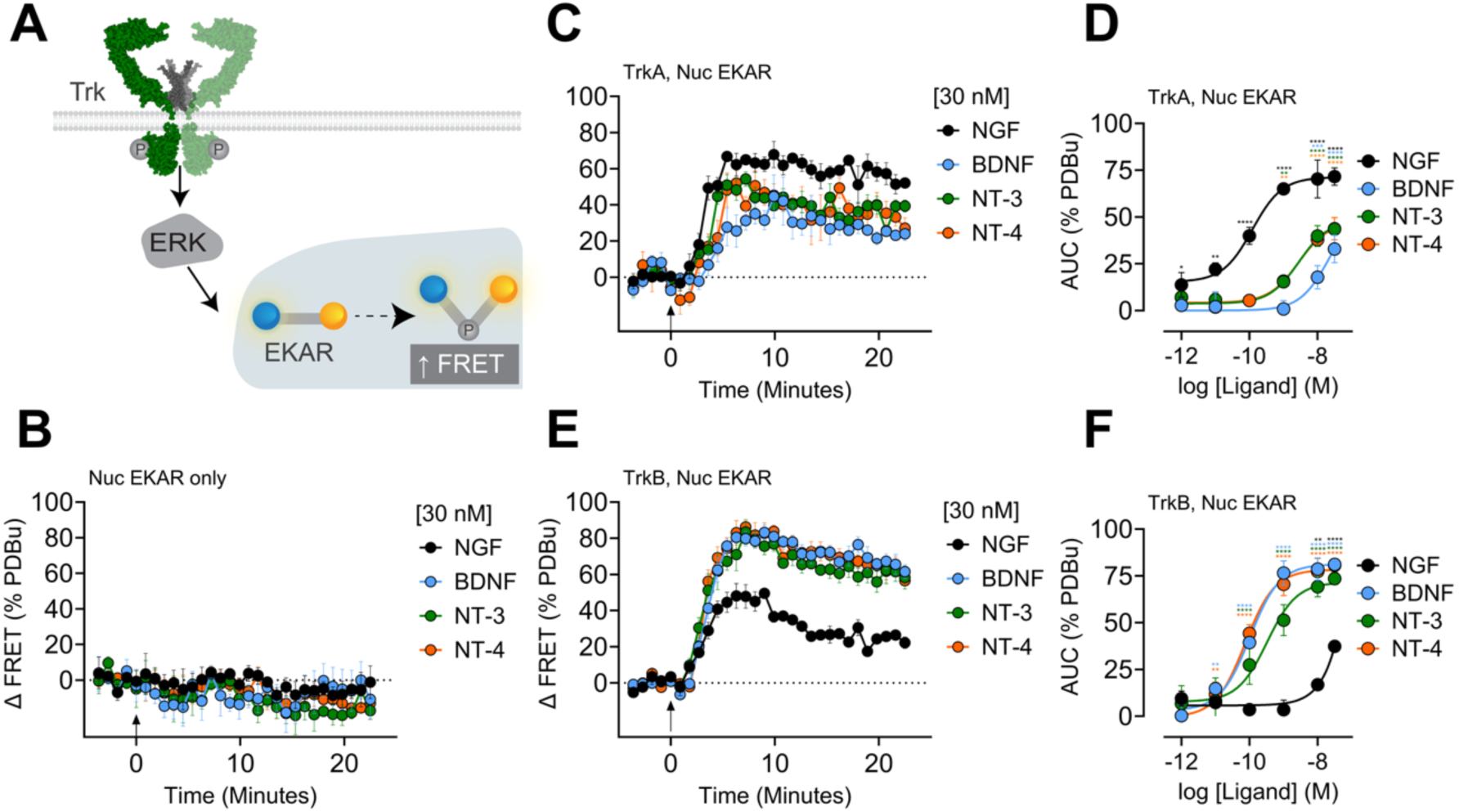
TrkA and TrkB have comparable ERK signalling. (A) Schematic outlining the nuclear EKAR fluorescence resonance energy transfer (FRET) reporter. (B) In HEK293T cells transfected with the EKAR reporter only, ERK signalling was measured in response to 30 nM of each neurotrophic growth factor, measured over 25 minutes (C,E) ERK signalling in TrkA- and TrkB-expressing HEK293T cells in response to 30 nM neurotrophic growth factor. Data expressed as change in FRET ratio over 25 minutes. (D,F) Concentration-response data obtained from the corresponding kinetic graph (Fig.1C,E) calculated as area under the curve (AUC). Data expressed as a percentage of vehicle (0%) and phorbol 12,13-dibutyrate (PDBu, 10 µM) (100%). Mean ± SEM, from 4-5 independent experiments with triplicate wells. Derived potency values and kinetic values are provided in Table 1. D,F. Ordinary two-way ANOVA with Dunnett’s multiple comparisons test versus vehicle. Statistical significance denoted as p<0.05 (*), p<0.01 (**), p<0.001 (***), and p<0.0001 (****).

**Table 1:** Summary of potency (pEC_50_) and kinetic (k_obs_) values derived for Trk nuclear ERK signalling. . Data are presented as mean ± SEM from 4-5 independent replicates with triplicate wells. While the pEC_50_ values are determined from the area under the curve, kinetic K_obs_ values were derived from between 2 & 15 minutes of the nuclear ERK signalling assay in response to 30 nM ligand. ‘Not determined’ (ND) indicates the response was insufficient to generate a pEC_50_ or K_obs_ value. Statistical analysis was performed comparing all ligands for each receptor against the canonical pairing (i.e TrkA/NGF vs TrkA/BDNF, TrkA/NT-3, and TrkA/NT-4), with significance denoted *p*<0.05 (*), *p*<0.01 (**), *p*<0.001 (***), and *p*<0.0001 (****); or performed across NT-3/receptor pairing, compared to TrkC_ΔKID_/NT-3 (i.e TrkC_ΔKID_/NT-3 vs TrkA/NT-3, TrkB/NT-3, TrkC_KID_/NT-3) with significance denoted as *p*<0.05 (^†^), *p*<0.01 (^††^), *p*<0.001 (^†††^), and *p*<0.0001 (^††††^).

|  | **TrkA** |  | **TrkB** |  | **TrkC KID** |  | **TrkC ΔKID** |  |
|---|---|---|---|---|---|---|---|---|
|  | pEC<sub>50</sub> | K<sub>obs</sub> (min<sup>-1</sup>) | pEC<sub>50</sub> | K<sub>obs</sub> (min<sup>-1</sup>) | pEC<sub>50</sub> | K<sub>obs</sub> (min<sup>-1</sup>) | pEC<sub>50</sub> | K<sub>obs</sub> (min<sup>-1</sup>) |
| NGF | 9.91 ± 0.09 | 2.54 ± 1.53 | ND | 1.22 ± 0.27 | ND | ND | ND | ND |
| BDNF | ND | ND | 9.95 ± 0.10 | 0.77 ± 0.21 | ND | ND | 8.26 ± 0.13** | 0.18 ± 0.02* |
| NT-3 | 8.45 ± 0.14****<br>†† | 1.14 ± 0.33<br>† | 9.48 ± 0.18 | 1.32 ± 0.26*<br>† | ND | 0.14 ± 0.06 | 9.55 ± 0.29 | 0.30 ± 0.03 |
| NT-4 | 8.50 ± 0.17**** | 0.90 ± 0.25 | 10.20 ± 0.10 | 1.24 ± 0.09 | ND | ND | ND | ND |

Cells were then systematically transfected with full-length, human TrkA or TrkB. Stimulation with NGF or BDNF respectively, their canonical ligands, induced robust ERK signalling within 10 minutes, reflected by a large increase in the FRET ratio (*Figure 1C, E*). This was observed in response to canonical and non-canonical ligands. While stimulation with NGF induced the largest amplitude in ERK signalling for TrkA, all ligands could induce a response in TrkA and TrkB to varying extents (*Figure 1C, E*). The real-time nature of this FRET-based ERK assay enabled kinetic parameters to be derived (*K_obs_; Table 1*). ERK signalling was then monitored in response to increasing ligand concentrations, plotted on concentration-response curves (*Figure 1D, F)*, from which the potency of ligand-induced nuclear ERK signalling was derived (*pEC_50_; Table 1*). ERK signalling downstream of TrkA was observed in response to all neurotrophic growth factors, with NGF producing the strongest and most potent change in the FRET ratio (EC_50_ ∼0.12 nM; *Table 1*). For both BDNF-induced TrkA and NGF-induced TrkB signalling, there was a modest increase in FRET at concentrations exceeding 10 nM, however no EC_50_ could be calculated from this concentration range (*Figure 1D, F*). TrkB then showed induced a strong ERK response when stimulated with BDNF, NT-3, and NT-4 all inducing concentration-dependent ERK activation (*Figure 1E, F*). While each neurotrophic factor showed high potency at TrkB, NT-4 was the most potent at inducing ERK signalling *via* TrkB (EC_50_ ∼0.06 nM; *Table 1*).

### Minimal ERK signalling in response to TrkC_KID_

TrkC can undergo alternative splicing to generate a 14 amino acid insert in the kinase domain adjacent to autophosphorylation sites in the activation loop (TrkC_KID_) (*Figure 2A*). While this 839aa TrkC_KID_ is the canonical isoform on UniProt, there is an alternative 825aa isoform of TrkC^24^ excluding these exon 16-encoded residues (denoted TrkC_ΔKID_ here). Here we take advantage of this isolated system to compare the real-time ERK signalling profile of both these isoforms utilising the nuclear EKAR reporter. The two TrkC isoforms showed markedly divergent ERK signalling profiles. TrkC_KID_ had a limited ERK response to neurotrophic ligands, where NT-3 elicited a weak increase in the nuclear ERK response at 30 nM reaching a peak of ∼35% of positive control PDBu (*Figure 2B,C*). Conversely, TrkC_ΔKID_ mediated robust ligand-induced ERK signalling comparable to TrkA and TrkB, when stimulated with NT-3 and BDNF (*Figure2E, F*). Responses to NT-3 and BDNF were both concentration-dependent (*Figure 2F*), with NT-3 being significantly more potent and faster at TrkC_ΔKID_ (EC_50_ ∼5.50 nM) compared to TrkC_KID_ (*Table 1*). BDNF induced an ERK response at TrkC_ΔKID_, however only activating the receptor at concentrations exceeding 1 nM. In terms of kinetics, TrkC_KID_ mediated ERK activation was particularly delayed with an ERK response only beginning after ∼5 minutes with a peak after ∼10 minutes (*Figure 2A)*, even at the highest NT-3 concentration at the top of the concentration-response curve. This was slower than ERK signalling in response to TrkC_ΔKID_, which all saw an increase in ERK activation after ∼3 minutes and peaking within ∼10 minutes after addition of their NT-3 ligand (*Figure 2E*) and BDNF (*Figure 2F*). TrkC_KID_ ERK activation via NT-3 was also significantly slower than TrkA/NGF and TrkB/BDNF (*K_obs_; Table 1*), demonstrating TrkC_KID_ uniquely slow/delayed kinetics in terms of ERK signalling. Given the striking difference in ERK signalling between TrkC_KID_ and TrkC_ΔKID_, we next investigated at which step in this receptor activation cascade this divergence originates.

**Figure 2.**
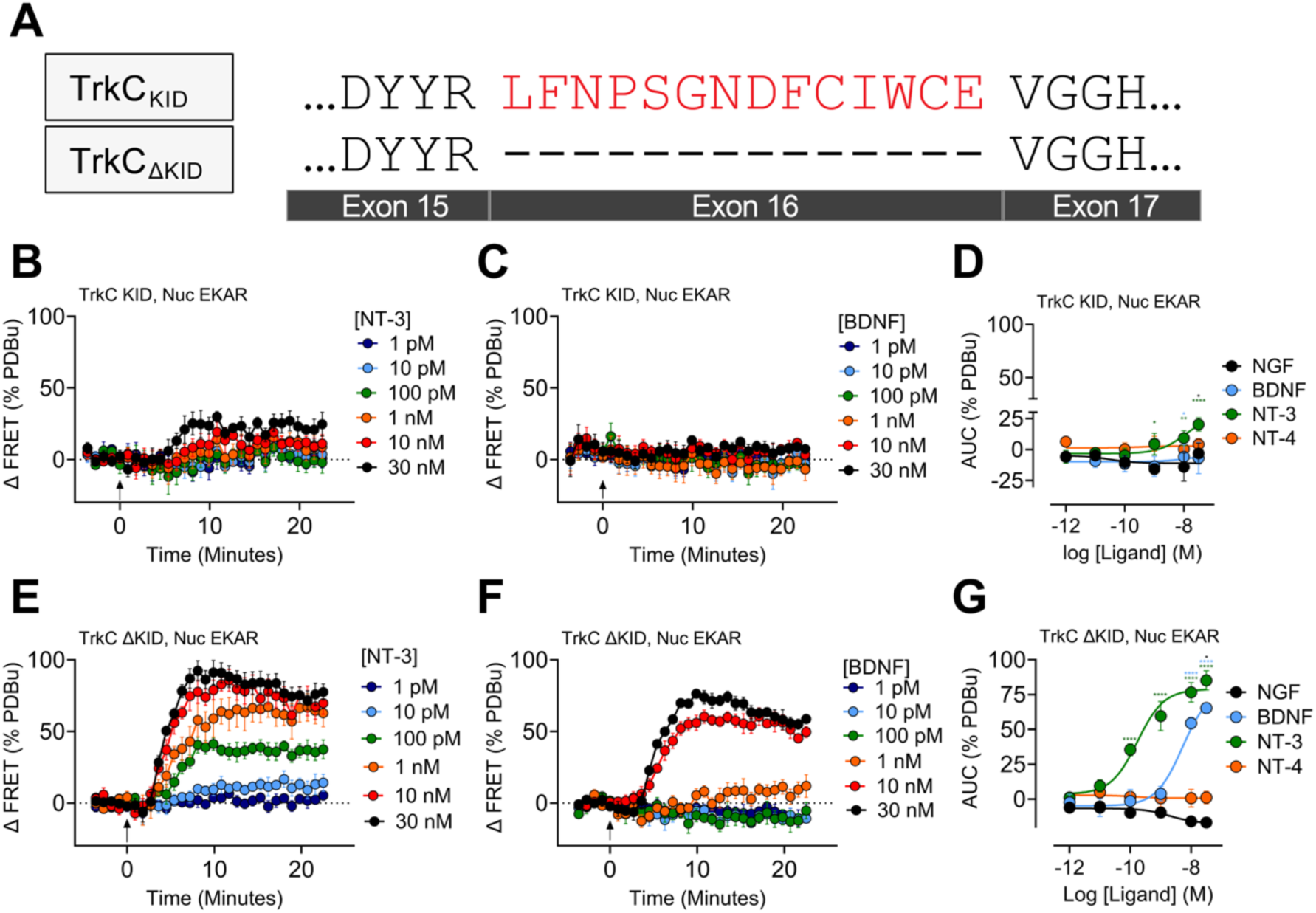
TrkC_KID_ had slower and reduced ERK signalling compared to TrkC_ΔKID_. (A) Schematic demonstrating alternative splicing of exon 16, leading to a 14 amino acid kinase insert domain (KID). (B,C,E,F) TrkC_KID_ and TrkC_ΔKID_ ERK signalling was measured in response to 30 nM of BDNF or NT-3. Data expressed as change in FRET ratio over 25 minutes. (D,G) Concentration-response data obtained from the corresponding kinetic graph (Fig.2C,F) calculated as area under the (AUC). Data expressed as a percentage of vehicle (0%) and Phorbol 12,13-dibutyrate (PDBu, 10 µM) (100%). Mean ± SEM, from 5 independent experiments with triplicate wells. Derived potency values and kinetic values are provided in Table 1. Significance (D,G) was determined using ordinary two-way ANOVA with Dunnett’s multiple comparisons test versus vehicle. Statistical significance denoted as p<0.05 (*), p<0.01 (**), p<0.001 (***), and p<0.0001 (****).

### Development of a Trk dimerisation assay

To investigate Trk receptor dimerisation, we developed a BRET-based assay to measure the proximity of Trk monomers following ligand stimulation. To facilitate this, a NanoLuciferase (NanoLuc, luminescent donor) and SnapTag (fluorescent acceptor) were tagged on the N-terminus of each monomer (*Figure 3A*). We first assessed the effect of canonical ligands for each Trk subtypes on their real-time dimerisation in response to increasing concentrations of neurotrophin (*Figure 3B-E*). At their cognate Trk receptor, each ligand induced a robust increase in the BRET ratio in a concentration-dependent manner. NGF-induced TrkA dimerisation and both BDNF- and NT-4-induced TrkB dimerisation demonstrated similar changes in the BRET ratio (*Figure 3B-C*). TrkC_KID_ and TrkC_ΔKID_, on the other hand, had a more pronounced increase in the BRET ratio in response to NT-3 compared to TrkA or TrkB (*Figure 3)*. Receptor expression levels were comparable across constructs, as determined by similar basal NanoLuc emission (luminescence), and pre-ligand BRET ratios were similar across Trk receptor subtypes (*Supplementary Figure 1*). Due to these distinct canonical ligands, it is unknown whether this effect is driven by NT-3 or by an intrinsic property of the TrkC receptor.

**Figure 3.**
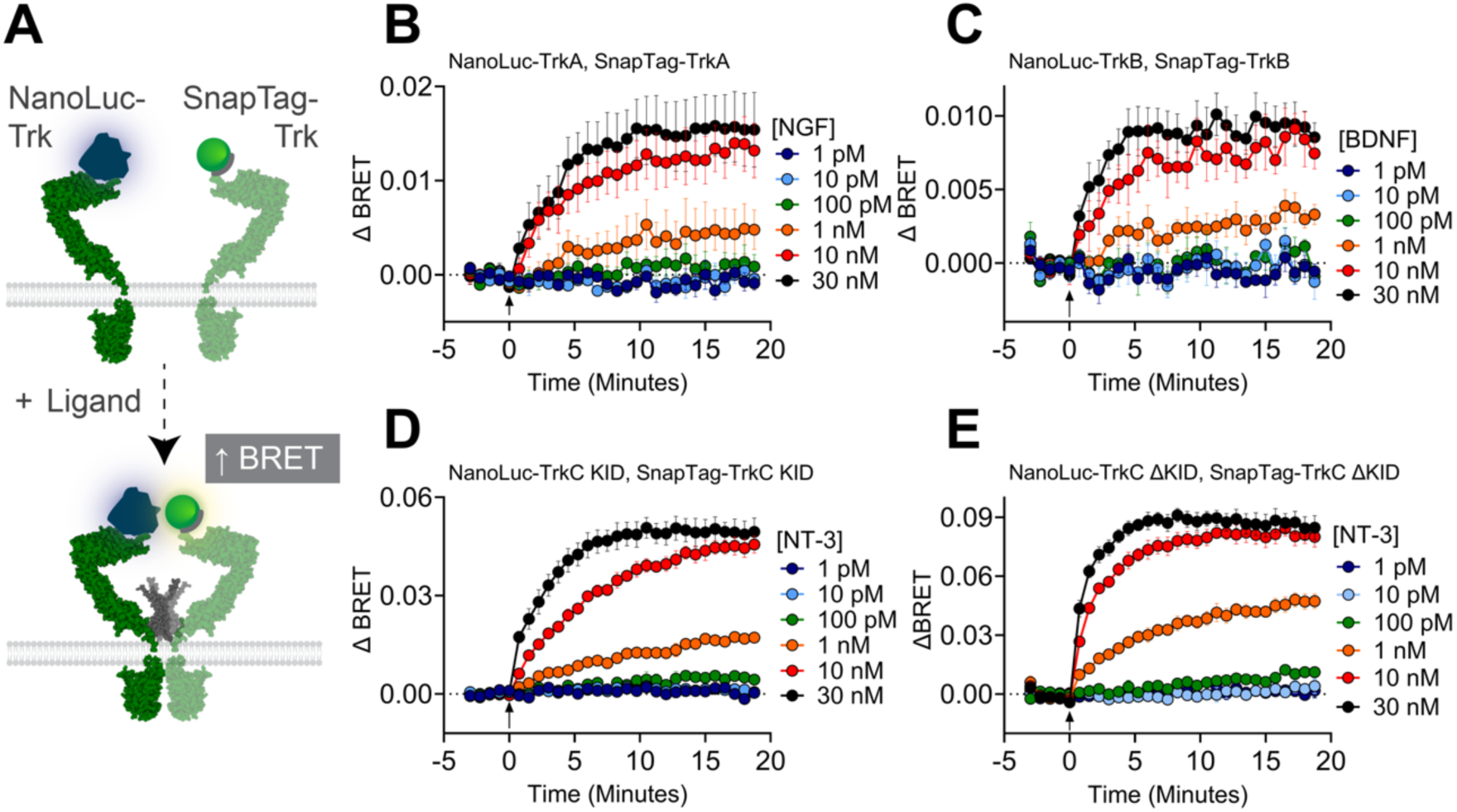
**Developing a real-time assay monitoring Trk dimerisation in response to canonical neurotrophin ligands**. (A) Schematic of the dimerisation assay measuring bioluminescence resonance energy transfer (BRET) between the luminescent donor enzyme (NanoLuc) and fluorescent acceptor (SnapTag-AlexaFluor488). (B-E) HEK293T cells expressing NanoLuc- and SnapTag-labelled TrkA (B), TrkB (C), TrkC_KID_ (D), or TrkC_ΔKID_ (E) were stimulated with increasing growth factor concentrations at T=0 (arrow). Data expressed as the change in BRET ratio over time. Mean ± SEM, from 4-5 independent experiments with triplicate wells.

### TrkC demonstrates atypical dimerisation compared to TrkA and TrkB

To probe whether these distinctions in TrkC/NT-3 are ligand-associated or receptor-associated, we monitored Trk receptor dimerisation in response to both canonical and non-canonical ligands. Dimerisation of TrkA was induced by NGF, and to a lesser extent BDNF, NT-3, and NT-4 (*Figure 4A*). Due to this concentration-dependent response, the BRET ratio following ∼20-minute stimulation was plotted as a concentration-response curve (*Figure 4B*). Quantifying the potency of neurotrophin-induced TrkA dimerisation (pEC_50_, *Table 2*), NGF had an EC_50_ ∼1.3 nM while the potency of TrkA dimerisation was comparable for NT-3 and NT-4. As well as deriving the potency for each neurotrophin, we used these real-time methods to quantify the kinetic parameters of neurotrophin-induced dimerisation (K_obs_; *Table 2)*. TrkB dimerisation was then observed in response to BDNF, NT-3, and NT-4 (*Figure 4C*). This rapid and robust increase in BRET ratio was concentration-dependent (*Figure 4D*), NT-4 was the most potent at inducing dimerisation of TrkB dimerisation (EC_50_ 0.6 nM). BDNF and NT-3 provided similar potencies (EC_50_ 1.9 - 2.6 nM; *Table 2*). NGF, however, did not induce any significant change in BRET ratio throughout the duration of the experiment.

**Figure 4:**
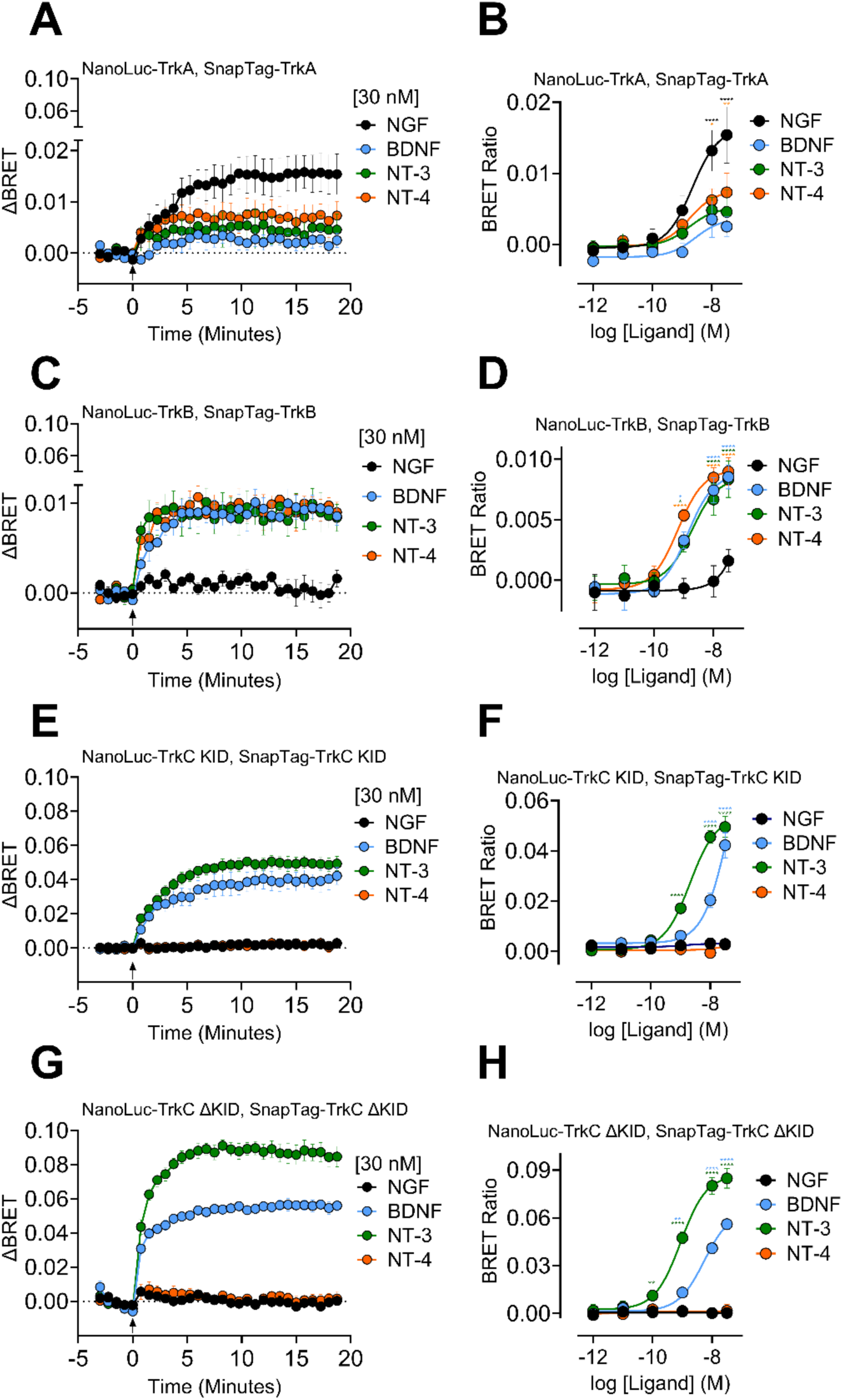
Concentration-dependent dimerisation of TrkA, TrkB, TrkC_KID_, and TrkC_ΔKID_ across the neurotrophin family. (A,C,E,G) Dimerisation between TrkA, TrkB, TrkC KID, or TrkC ΔKID for all four neurotrophins (30 nM), including the top concentration of canonical ligand from Fig.3. (B,D,F,H) Concentration-response across a range of concentrations plotting the baseline-corrected BRET ratio from ∼20 minutes. Mean ± SEM, from 4-5 independent experiments with triplicate wells. Potency values (pEC_50_, Table 2) and kinetic parameters (k_obs_, Table 1) were fit using a non-linear regression. Significance (B,D,F,H) was determined using ordinary two-way ANOVA with Dunnett’s multiple comparisons test versus vehicle. Statistical significance denoted as p<0.05 (*), p<0.01 (**), p<0.001 (***), and p<0.0001 (****).

**Table 2:** Summary of potency (pEC_50_) and kinetic (k_obs_) values derived for Trk dimerisation. . Data are presented as mean ± SEM from 5 independent replicates with triplicate wells. K_obs_ values were derived from Trk dimerisation in response to 30 nM ligand. ‘Not determined’ (ND) indicates the response was insufficient to generate a pEC_50_ or K_obs_ value. Statistical analysis was performed comparing all ligands for each receptors against the canonical pairing (i.e TrkA/NGF vs TrkA/BDNF, TrkA/NT-3, and TrkA/NT-4), with significance denoted *p*<0.05 (*), *p*<0.01 (**), *p*<0.001 (***), and *p*<0.0001 (****); or performed across NT-3/receptors pairing, compared to TrkC_ΔKID_/NT-3 (i.e TrkC_ΔKID_/NT-3 vs TrkA/NT-3, TrkB/NT-3, TrkC_KID_/NT-3) with significance denoted as *p*<0.05 (^†^), *p*<0.01 (^††^), *p*<0.001 (^†††^), and *p*<0.0001 (^††††^).

|  | **TrkA** |  | **TrkB** |  | **TrkC KID** |  | **TrkC ΔKID** |  |
|---|---|---|---|---|---|---|---|---|
|  | pEC<sub>50</sub> | K<sub>obs</sub> (min<sup>-1</sup>) | pEC<sub>50</sub> | K<sub>obs</sub> (min<sup>-1</sup>) | pEC<sub>50</sub> | K<sub>obs</sub> (min<sup>-1</sup>) | pEC<sub>50</sub> | K<sub>obs</sub> (min<sup>-1</sup>) |
| NGF | 8.88 ± 0.44 | 0.33 ± 0.04 | ND | ND | ND | ND | ND | ND |
| BDNF | ND | ND | 8.72 ± 0.24 | 0.56 ± 0.08 | ND | ND | 8.29 ± 0.08** | 0.89 ± 0.16 |
| NT-3 | 8.99 ± 0.56 | 1.24 ± 0.57 | 8.59 ± 0.18 | 2.02 ± 0.64* | 8.70 ± 0.07 | 0.39 ± 0.03 | 9.06 ± 0.14 | 0.84 ± 0.12 |
| NT-4 | 8.58 ± 0.34 | 0.86 ± 0.22 | 9.22 ± 0.10 | 0.86 ± 0.10 | ND | ND | ND | ND |

A large change in BRET upon TrkC_KID_ and TrkC_ΔKID_ dimerisation was observed in response to both NT-3 and BDNF (*Figure 4E,G*). This was concentration-dependent (*Figure 4F, H*), where the potency for NT-3 for TrkC_KID_ (EC_50_ ∼2 nM) and TrkC_ΔKID_ (EC_50_ ∼0.87 nM) indicates that the KID does not impact the enhanced TrkC dimerisation when measured using tags on the N-terminus. In both cases, NT-3 induced a more potent response compared to BDNF with respect to dimerisation (*Table 2*). Both NGF and NT-4, on the other hand, failed to induce any change in BRET ratio for TrkC_KID_ and TrkC_ΔKID_ (*Figure 4F,H*). As the dimerisation response for both TrkC_KID_ and TrkC_ΔKID_ were distinct from TrkA or TrkB, we assessed whether this was linked to differences in receptor trafficking.

### Comparing real-time trafficking across the Trk family

We set out to compare the real-time trafficking of Trk receptors away from the plasma membrane following ligand stimulation across the Trk family. To achieve this, a distinct BRET-based assay was established tagging the C-terminus with Rluc8 and using a RGFP-CAAX as a plasma membrane marker (*Figure 5A*). Upon ligand stimulation, receptor internalisation leads to trafficking away from the plasma membrane and the RGFP-CAAX marker, meaning a decrease in BRET ratio. In response to increasing concentrations of its canonical ligand, there was trafficking of Trk receptors away from the plasma membrane marker with a sustained reduction in BRET signal (*Figure 5B-E*). Concentration-response curves (*Figure 6B,D,F,H*) were generated from baseline-corrected BRET ratio at ∼20-minutes of the corresponding kinetic graphs (*Figure 6A,C,E,G*). From these data, the potency of ligand-induced trafficking (pEC_50_; *Table 3*), and kinetic parameters (*K_obs_; Table 3*) were derived to quantify difference across Trk receptors. TrkA trafficking was most potently induced by NGF (EC_50_ 0.95 nM; *Table 3*), with NT-3 and NT-4 eliciting less potent responses (EC_50_ 14.8 nM and 6.92 nM respectively; *Table 3*). BDNF was unable to produce comparative TrkA trafficking (*Figure 6A*). TrkB trafficking was unique in the sense that all ligands could induce a measurable change in BRET, with BDNF being the most potent (EC_50_ 2.34 nM; *Table 3*) and NGF being the least potent (EC_50_ 8.91 nM) whilst also only inducing a partial trafficking response compared to the other growth factors (*Figure 6D*). NT-3 and NT-4 induced a similar TrkB trafficking response with potencies of 3.31 nM and 3.39 nM respectively (*Table 3*).

**Figure 5.**
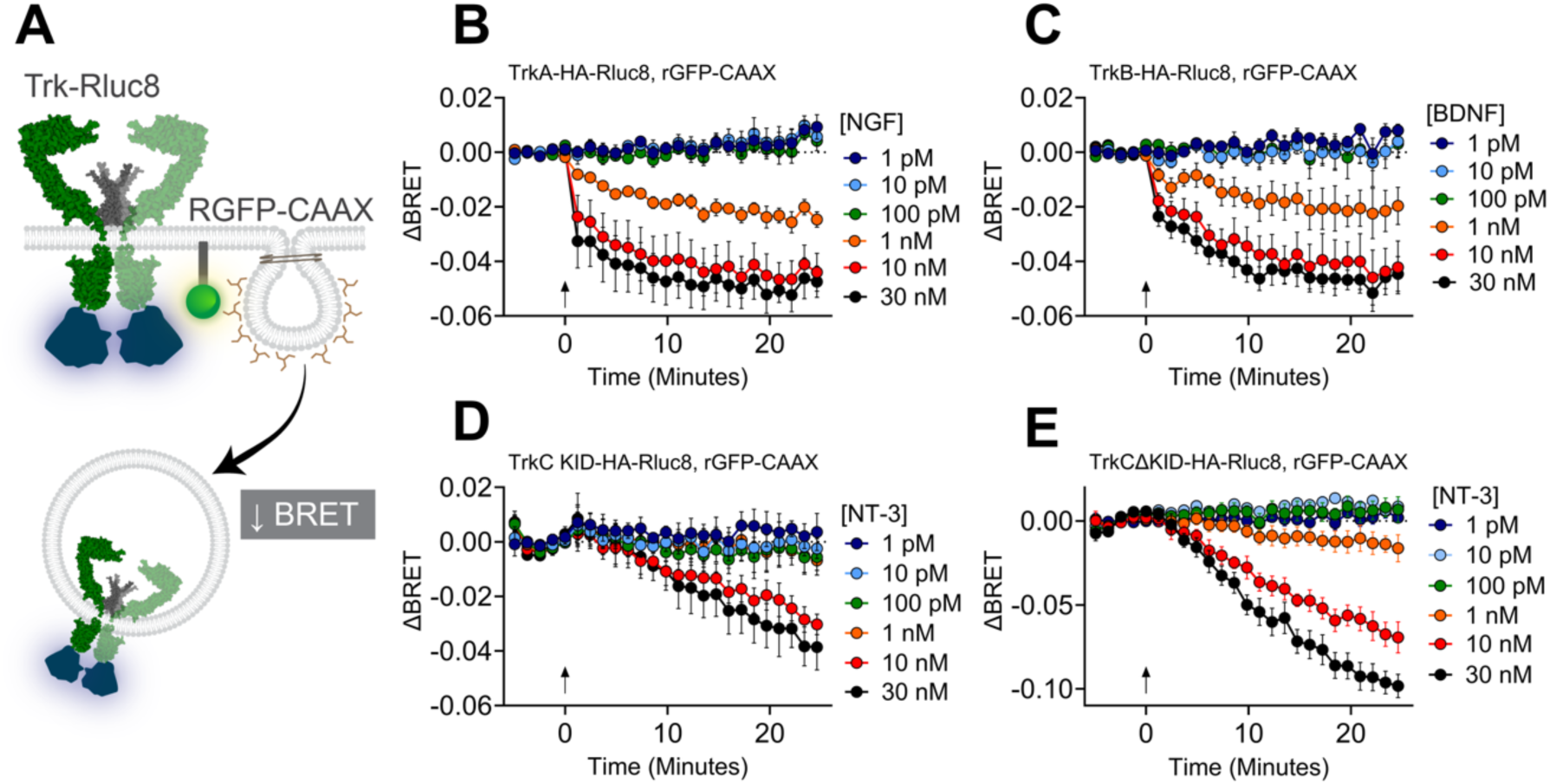
Monitoring Trk trafficking in real-time in response to canonical neurotrophin ligands. (A) Schematic of the Trk trafficking assay measuring BRET between Trk-HA-Rluc8 (bioluminescent donor) and RGFP-CAAX (fluorescent acceptor). (B-E) Trafficking interactions away from the plasma membrane in response to canonical growth factors. Data are expressed as change in BRET ratio over 25 minutes n=5. Ligand (1 pM - 30 nM) was added at T=0 (arrow). Data points represent mean ± SEM, from 5 independent experiments with triplicate wells.

**Figure 6.**
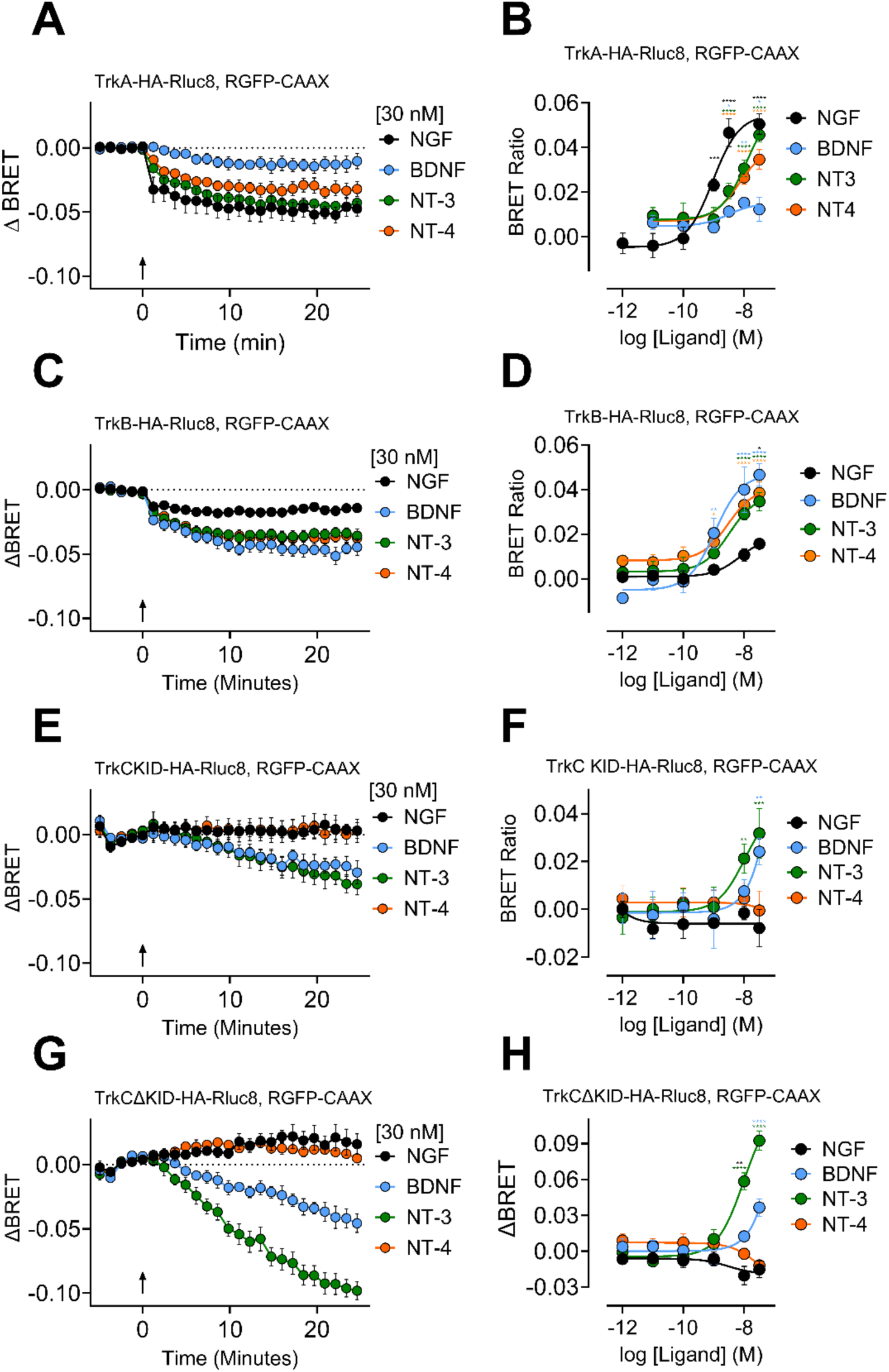
TrkC_KID_ had reduced trafficking away from the plasma membrane marker compared to TrkA, TrkB, or TrkC_ΔKID_. (A,C,E,G) Trafficking in response to 30 nM of each neurotropic growth factors. Data are expressed as change in BRET ratio over 25 minutes. (B,D,F,H) Concentration–response curves were generated from the baseline-corrected BRET ratio at ≈20 min of the corresponding kinetic graphs (Fig. 5B–E) and inverted to reflect growth factor–induced receptor endocytosis. Mean ± SEM, from 5 independent experiments with triplicate wells. Derived potency and kinetic values are provided in Table 3. Significance (B,D,F,H) was determined using ordinary two-way ANOVA with Dunnett’s multiple comparisons test versus vehicle. Statistical significance denoted as p<0.05 (*), p<0.01 (**), p<0.001 (***), and p<0.0001(****).

**Table 3:**
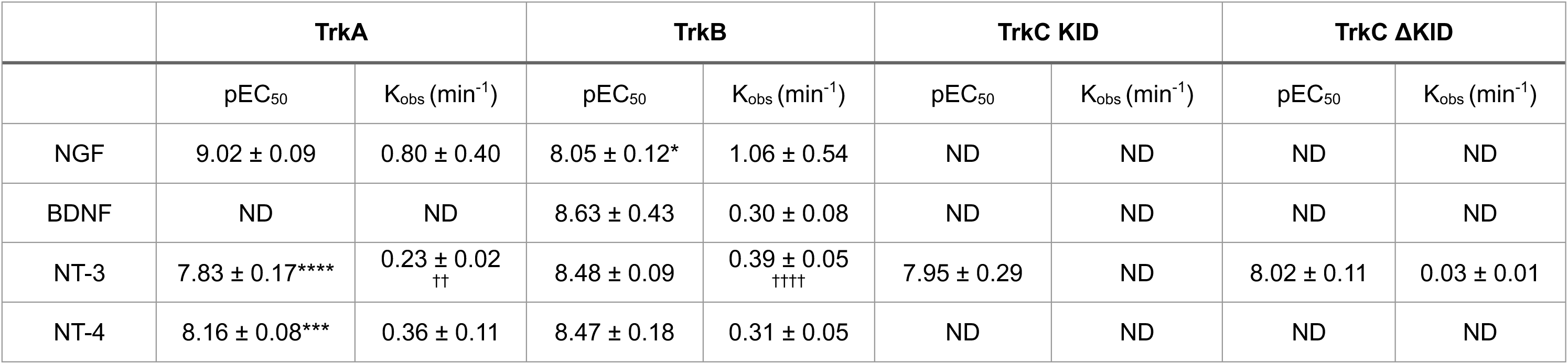
Summary of potency (pEC_50_) and kinetic (k_obs_) values derived for Trk trafficking. . Data are presented as mean ± SEM from 4-5 independent replicates with triplicate wells. K_obs_ values were derived from Trk trafficking in response to 30 nM ligand. ‘Not determined’ (ND) indicates the response was insufficient to generate a pEC_50_ or K_obs_ value. Statistical analysis was performed comparing all ligands for each receptors against the canonical pairing (i.e TrkA/NGF vs TrkA/BDNF, TrkA/NT-3, and TrkA/NT-4), with significance denoted *p*<0.05 (*), *p*<0.01 (**), *p*<0.001 (***), and *p*<0.0001 (****); or performed across NT-3/receptor pairing, compared to TrkC_ΔKID_/NT-3 (i.e TrkC_ΔKID_/NT-3 vs TrkA/NT-3, TrkB/NT-3, TrkC_KID_/NT-3) with significance denoted as *p*<0.05 (^†^), *p*<0.01 (^††^), *p*<0.001 (^†††^), and *p*<0.0001 (^††††^).

|  | **TrkA** |  | **TrkB** |  | **TrkC KID** |  | **TrkC ΔKID** |  |
|---|---|---|---|---|---|---|---|---|
|  | pEC<sub>50</sub> | K<sub>obs</sub> (min<sup>-1</sup>) | pEC<sub>50</sub> | K<sub>obs</sub> (min<sup>-1</sup>) | pEC<sub>50</sub> | K<sub>obs</sub> (min<sup>-1</sup>) | pEC<sub>50</sub> | K<sub>obs</sub> (min<sup>-1</sup>) |
| NGF | 9.02 ± 0.09 | 0.80 ± 0.40 | 8.05 ± 0.12* | 1.06 ± 0.54 | ND | ND | ND | ND |
| BDNF | ND | ND | 8.63 ± 0.43 | 0.30 ± 0.08 | ND | ND | ND | ND |
| NT-3 | 7.83 ± 0.17**** | 0.23 ± 0.02 †† | 8.48 ± 0.09 | 0.39 ± 0.05 †††† | 7.95 ± 0.29 | ND | 8.02 ± 0.11 | 0.03 ± 0.01 |
| NT-4 | 8.16 ± 0.08*** | 0.36 ± 0.11 | 8.47 ± 0.18 | 0.31 ± 0.05 | ND | ND | ND | ND |

The TrkC isoforms produced vastly different responses in trafficking assay compared to TrkA and TrkB. Only NT-3 induced a noticeable reduction in BRET in a concentration-dependent manner (*Figure 6E; Table 3;* EC_50_ ∼11.22 nM for TrkC_KID_ and ∼9.55 nM for TrkC_ΔKID_). BDNF was able to induce TrkC_KID_ and TrkC_ΔKID_ trafficking, but only at 30 nM, meaning an pEC_50_ couldn’t be calculated for BDNF. NGF and NT-4 failed to induce any trafficking in either TrkC isoform (*Figure 6E,F,G,H*). Most notably, TrkC_KID_ and TrkC_ΔKID_ had noticeably slower internalisation compared to TrkA/NGF and TrkB/BDNF (*Table 3;* TrkA K_obs_ 0.23 min^-1^, TrkB K_obs_ 0.39 min^-1^, and TrkC_ΔKID_ K_obs_ 0.03 min^-1^). This demonstrates that the receptor is unique in its dynamics, even irrespective of the removal of the KID and upon restoration of TrkC signalling. Overall, both TrkC isoforms demonstrated unique receptor trafficking dynamics.

### Distinct TrkC dimerisation and trafficking is independent of cleavage mutations

As TrkC has been proposed as a dependence receptor through cleavage of the intracellular receptor^15^, we generated caspase-resistant mutants at the two major cleavage sites by substituting aspartic acid with asparagine (D495N and D641N) in TrkC_KID_ (*Figure 7A*) to interrogate whether this was responsible for distinct dimerisation and trafficking compared to TrkA/TrkB. Receptor expression levels and surface expression were first assessed across TrkC variants by measuring baseline luminescence (NanoLuc and Rluc8; *Supplementary Figure 2*), as well as the extent of TrkC expressed at the cell surface (*Supplementary Figure 2*). TrkC_KID_ (D495N), TrkC_KID_ (D641N), and TrkC_KID_ (WT) all demonstrated similar baseline NanoLuc luminescence and pre-ligand baseline BRET (*Supplementary Figure 2*), confirming that receptor expression levels were comparable between mutants. Both TrkC_KID_ (D495N) and TrkC_KID_ (D641N) produced a similar dimerisation response compared to TrkC_KID_ (WT) (*Figure 7B*). Both mutations also showed no distinct in receptor trafficking relative to wild type, with the D641N mutant displaying a modestly weaker response (*Figure 7C*), however the unique TrkC kinetic profile TrkC remaining comparable across both mutants with respect to TrkC trafficking. While Rluc8-tagged receptor expression was similar, the constitutive amount of TrkC at the cell surface quantified by BRET between Rluc8 and rGFP is reduced in TrkC_ΔKID_ and TrkC_KID_ (D641N), although this did not impact the dynamics of receptor internalisation. Furthermore, ERK signalling of TrkC_KID_ (D495N) and TrkC_KID_ (D641N) remained very similar to TrkC_KID_ and achieved only ∼35% of maximal PDBu response (*Figure 7D*). This demonstrates that the unique spatiotemporal dynamics of TrkC are independent of its ability to be cleaved within the intracellular portion of the receptor.

**Figure 7.**
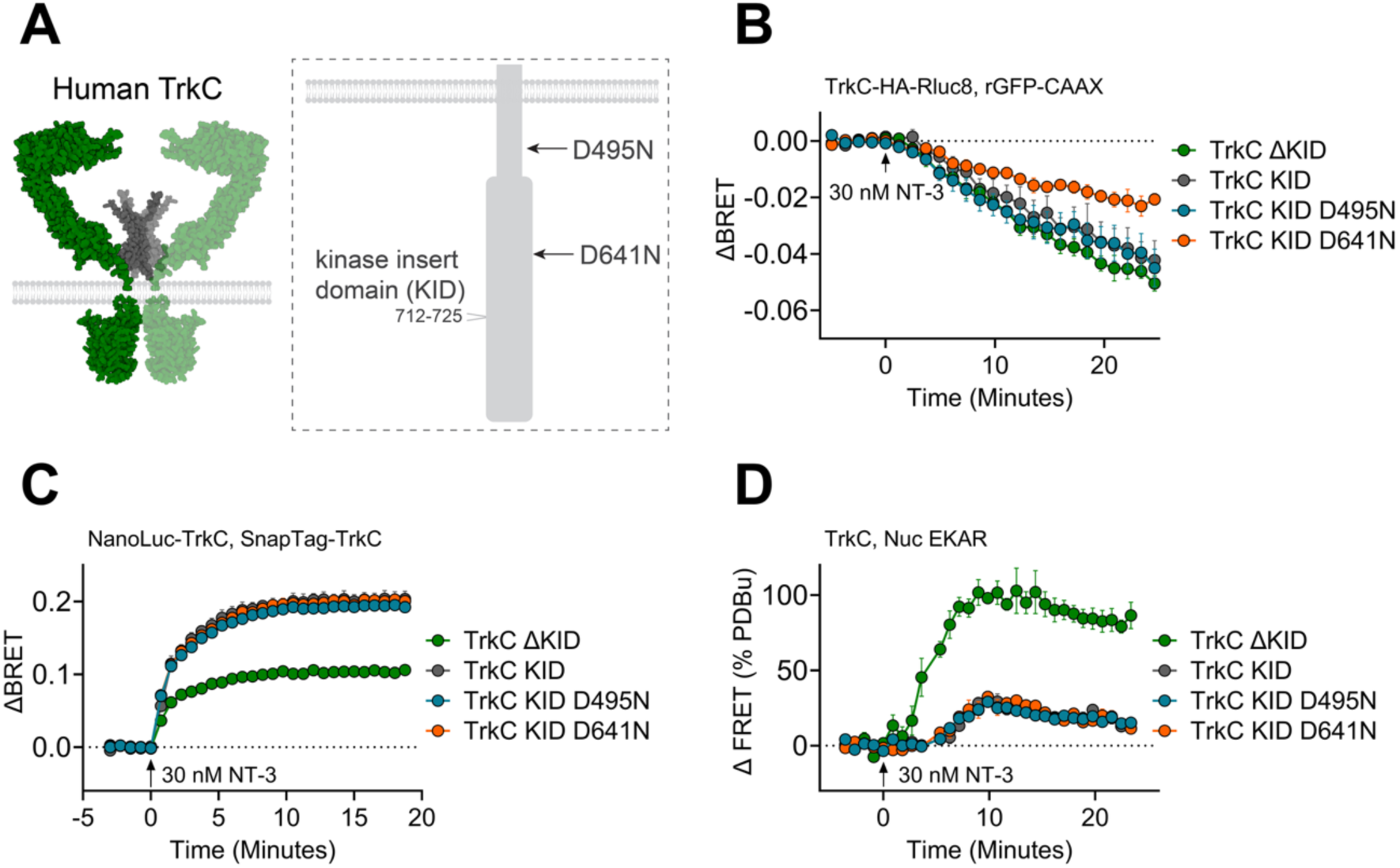
Caspase-resistant TrkC_KID_ mutations have no impact on receptor internalisation, dimerisation, or ERK signalling. (A) Schematic representation of TrkC mutations interfering with caspase cleavage (D495N and D641N). (B) Receptor trafficking from HEK293T cells expressing C-terminally Rluc8-tagged TrkC_ΔKID_, TrkC_KID_, or TrkC_KID_ (D495N) or TrkC_KID_ (D641N) by stimulating with 30 nM NT-3 at T=0. (C) HEK293T cells expressing NanoLuc- and SnapTag-labelled TrkC_ΔKID_, TrkC_KID_, TrkC_KID_ (D495N) or TrkC_KID_ (D641N) were stimulated 30 nM of NT-3 at T=0. (D) ERK signalling was measured in cells expressing TrkC_ΔKID_, TrkC_KID_, or TrkC_KID_ (D495N) or TrkC_KID_ (D641N), in response to 30 nM NT-3. Data expressed as the change in BRET ratio over time. Data is expressed as change in FRET ratio and expressed as a percentage of vehicle (0%) and PDBu (100%). Mean ± SEM, from 5 independent experiments with triplicate wells.

## Discussion

Our results provide novel insights into Trk receptor dynamics, revealing that two isoforms of TrkC exhibit distinct characteristics compared to TrkA and TrkB. TrkC was unique in that it exhibits enhanced receptor dimerisation, as well as having slower endocytosis dynamics, compared to TrkA and TrkB. This was identified using a BRET-based real-time assay at 37°C to measure how these specifically tagged human receptors dimerise or move away from the plasma membrane. While the TrkC isoform lacking the KID (TrkC_ΔKID_, 825aa) displayed ERK signalling profiles comparable to TrkA and TrkB, the KID-containing isoform (TrkC_KID_, 839aa) is characterised by reduced nuclear ERK signalling, monitored using a downstream FRET-based assay to measure real-time ERK signalling at the human receptor expressed in isolation.

While Trk receptors are canonically paired with specific neurotrophins, our findings provide evidence for cross-family ligand/receptors interactions extending beyond promiscuous NT-3^29^, with some potency values for ‘non-canonical’ pairings comparable to their canonical ligands. Both NT-4 and NT-3, for example, were capable of inducing TrkA ERK signalling, dimerisation, and trafficking, with a similar potency to NGF with respect to the dimerisation assay. Similarly, BDNF robustly stimulated TrkC signalling, dimerisation and trafficking. Whilst BDNF had a lower potency than NT-3 for TrkC, these signalling dynamics were comparable between BDNF and NT-3. These assays were confirmed to be selective, as seen with the pairings that did not induce ERK signalling, dimerisation or trafficking – such as NGF/TrkB or NT-4/TrkC. Intriguingly, BDNF elicited modest ERK activation *via* TrkA at higher ligand concentrations, despite limited dimerisation or trafficking. This may occur through indirect or low-affinity interactions amplified by downstream signalling.

Interestingly, both TrkC isoforms exhibited a considerably larger dimerisation response compared with TrkA and TrkB. Cleavage-resistant mutants (D495N and D641N) in the TrkC_KID_ receptor maintained this elevated dimerisation response, indicating that caspase cleavage at Asp495 and Asp641 does not prevent receptor dimerisation as measured at the N-terminus. Published studies investigating Trk receptor dimerisation suggested a population of TrkA and TrkB can exist as pre-formed, inactive dimers at the plasma membrane under resting conditions^30,31^, where ligand binding is proposed to induce a conformational re-arrangement within these pre-existing complexes, thereby enabling downstream signalling. Although the dynamics of TrkC was not assessed in these studies, the BRET ratio prior to neurotrophin stimulation was comparable between TrkA, TrkB and TrkC. It should be noted that the change in the BRET signal could also reflect a ligand-induced conformational change within a preformed dimer that brings donor and acceptor tags into closer proximity. FRET studies investigating C-terminal tagged receptors support this TrkC difference in dimerisation, showing a distinctive FRET change suggesting TrkC has a unique conformational re-arrangement compared to TrkA and TrkB^32^. Our study using full-length receptors therefore supports a similar model for TrkC, irrespective of whether TrkC_ΔKID_ or signalling-deficient TrkC_KID_ were studied. This could be attributed to enhanced oligomerisation or a distinct conformational change in comparison to TrkA and TrkB, however structural information is lacking for the extracellular domain of TrkC.

Although TrkC functions as a dependence receptor that undergoes caspase-dependent cleavage^15^, the preservation of robust dimerisation in both these cleavage-resistant TrkC mutants suggests that the extent of dimerisation occurs independently from the cleavage event. Neither the inhibitory 14aa of the KID, nor cleavage at Asp495 and Asp641, substantially impaired ligand-induced TrkC dimerisation. This indicates that TrkC_KID_ retains affinity for NT-3 ligand binding and the ability to form receptor complexes at the cell surface. One mechanistic possibility underpinning this is that TrkC_KID_ may function as a ligand sink, similarly to truncated Trk isoforms^22^. By sequestering NT-3 and only inducing ∼35% of the signalling output of its insert lacking counterpart (TrkC_ΔKID_), this allows neurons to fine-tune local neurotrophin availability in the cellular environment. As TrkC_ΔKID_ is known to be expressed within the nervous system alongside both TrkC_KID_ and TrkC.T1, it important to understand how this alternative splicing of TrkC is regulated, particularly given then implications for therapeutic targeting of Trk receptors.

In our study, the two TrkC isoforms displayed distinct trafficking dynamics compared to TrkA and TrkB. TrkC_ΔKID_ underwent robust ligand-induced internalisation away from the plasma membrane, ultimately reaching an extent comparable to TrkA and TrkB. However, its kinetics were significantly delayed relative to TrkA and TrkB, with internalisation beginning later and proceeding more slowly. TrkA and TrkB are well documented to internalise via clathrin-coated vesicles following ligand binding^17,19^. Once internalised, both receptors undergo retrograde transport in signalling endosomes that support prolonged activation of ERK and PI3K/AKT pathways ^33–35^. In contrast, there is evidence that TrkC trafficking follows a distinct route primarily to the Golgi apparatus rather than classical endosomes^36^. This may contribute to the modestly delayed kinetics observed for both TrkC isoforms. While kinase activity is thought to be required for efficient clathrin-mediated endocytosis^37,38^, given we do not see any distinct in ERK signalling kinetics between TrkA, TrkB or TrkC, this would not justify the significantly slower trafficking kinetics observed for TrkC.

We also observed notably reduced ERK signalling downstream of TrkC_KID_ activation, compared to TrkC_ΔKID_. This agrees with published literature, attributed to this 14 amino acid KID directly adjacent to the “YXXDYY motif” in the activation loop – a major site of autophosphorylation is inhibited by the KID^24^. This agrees with the markedly reduced ERK activation in previous studies investigating TrkC_KID_^23,24^. Using a real-time FRET-based EKAR sensor at 37°C, our study provides the first direct, side-by-side comparison of the temporal dynamics of ERK signalling between the two TrkC isoforms and the other Trk receptors under identical conditions. This approach therefore clarifies the impact of the KID on TrkC dynamics, revealing not only the profound signalling deficit caused by the insert but also its specific impact on both the magnitude and the kinetics of the ERK response. Furthermore, cleavage-resistant mutations (D495N and D641N) in the TrkC_KID_ receptor did not rescue ERK signalling, confirming that the signalling defect is driven by the KID insert itself and is independent of caspase cleavage at Asp495 and Asp641.

Despite the distinctions observed in both TrkC isoforms from TrkA/B in terms of dynamics regarding receptor trafficking and downstream ERK signalling, their selectivity was similar between TrkC_KID_ and TrkC_ΔKID_. Specifically looking at TrkC_ΔKID_, the signalling form, NGF and NT-4 failed to induce any ERK signalling, dimerisation, or receptor trafficking. This effect can also clearly be seen for TrkC_KID_ dimerisation, with NGF and NT-4 failing to induce any change in BRET. Overall, this suggests that signalling-deficient TrkC_KID_ and signalling-competent TrkC_ΔKID_ have a higher selectivity for NT-3 but can also respond to BDNF at high concentrations, thus with lower potency. These observations therefore support a more nuanced model of Trk signalling, in which neurotrophin-receptor interactions are not strictly exclusive but context-dependent, with ligand concentration, receptor expression, and cellular environment contributing to the observed signalling landscape of neurotrophins.

These proximity-based approaches enabled us to probe the real-time pharmacology of specified human Trk receptors for each stage of the receptor signalling cascade, with the ability to study full length Trk receptors in living cells. By performing all assays at 37°C, real-time receptor trafficking, dimerisation, and signalling are reflective of their physiological temperature, something often lost with other approaches despite temperature having a considerable impact on kinetics^39^. These approaches are also selective for tagged receptor populations, ensuring that BRET and FRET measurements reflect the dynamics of the defined Trk constructs only – and the specific isoform – allowing a high level of selectivity. This is especially crucial given the high degree of homology between the Trk receptor family. Moreover, the high-throughput, plate-based format is well-suited for screening novel Trk agonists, such as *de novo* NGF mimetics^40^ and TrkB-targeting antibodies such as ZEB85^41^, making these powerful tools for drug discovery with both spatial and temporal resolution.

In conclusion, TrkC displayed markedly divergent spatial and temporal dynamics compared to TrkA and TrkB. Both TrkC isoforms studied – signalling-deficient TrkC_KID_ and TrkC_ΔKID_ – demonstrated a particularly strong dimerisation and delayed trafficking compared to TrkA and TrkB, which remained distinct even upon mutation of caspase cleavage sites. While TrkC_KID_ had severely impaired ERK signalling that was similarly independent of caspase cleavage sites, the 825aa TrkC_ΔKID_ isoform had comparable ERK signalling to TrkA and TrkB. Given the continued and growing interest in targeting Trk receptors across a range of neurodegenerative diseases^8,9^ and psychiatric disorders^42^, these isoform-specific and Trk receptor-specific mechanisms provide critical molecular insights into how these receptors differ despite their high homology, guiding future efforts to selectively modulate Trk receptor signalling therapeutically.

## Acknowledgements

This work was supported by the Biotechnology and Biological Sciences Research Council Doctoral Training Programme [BB/T0083690/1] and the Academy of Medical Sciences Springboard Award [SBF0010\1058].

## Author Contributions

Conceptualization: CJP; Methodology: CJP, RJD; Investigation: RJD; Formal analysis: RJD; Data curation: RJD, CJP; Supervision: CJP, SJH; Writing - original draft: RJD; CJP; Writing - review and editing: all authors.

## Data Availability

The data supporting the findings of this study are available within the article and its Supplementary Information. Source data underlying the graphs presented are available from the corresponding author upon request.

## Supplementary Figure

**Supplementary Figure 1.**
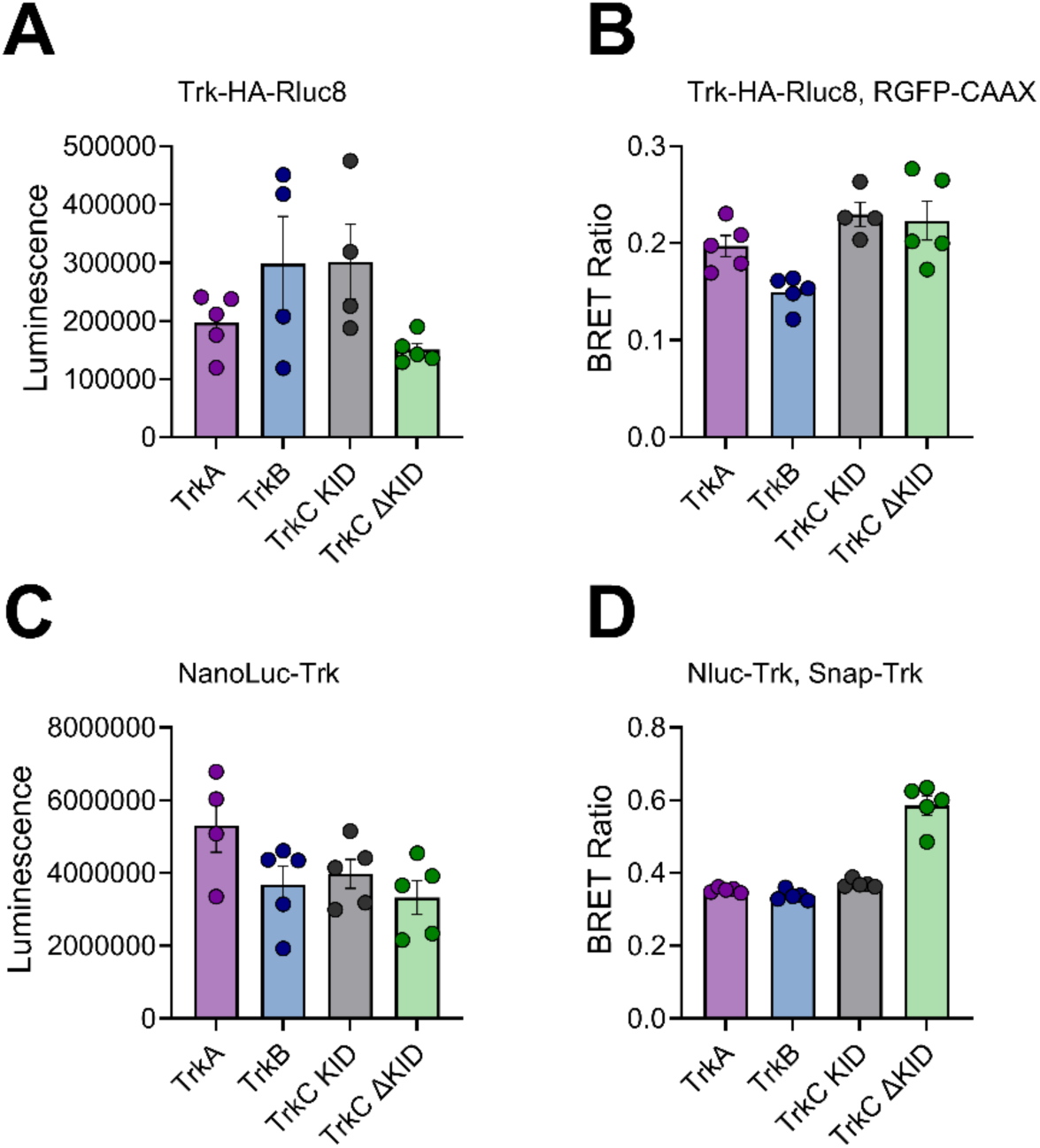
Luminescence emissions and pre-ligand addition BRET ratios for TrkA, TrkB, TrkC_KID_, and TrkC_ΔKID_. (A) Receptor expression (Rluc8 luminescence emissions) and (B) BRET ratio between C-terminal tagged luminescent donor (Rluc8) and fluorescent acceptor (RGFP-CAAX) in the absence of ligand. (C) Receptor expression (NanoLuc luminescence emissions) and (D) BRET ratio between N-terminal tagged luminescent donor (NanoLuc) and fluorescent acceptor (SnapTag-AlexaFluor488). Mean ± SEM, from 4-5 independent experiments with triplicate wells.

**Supplementary Figure 2.**
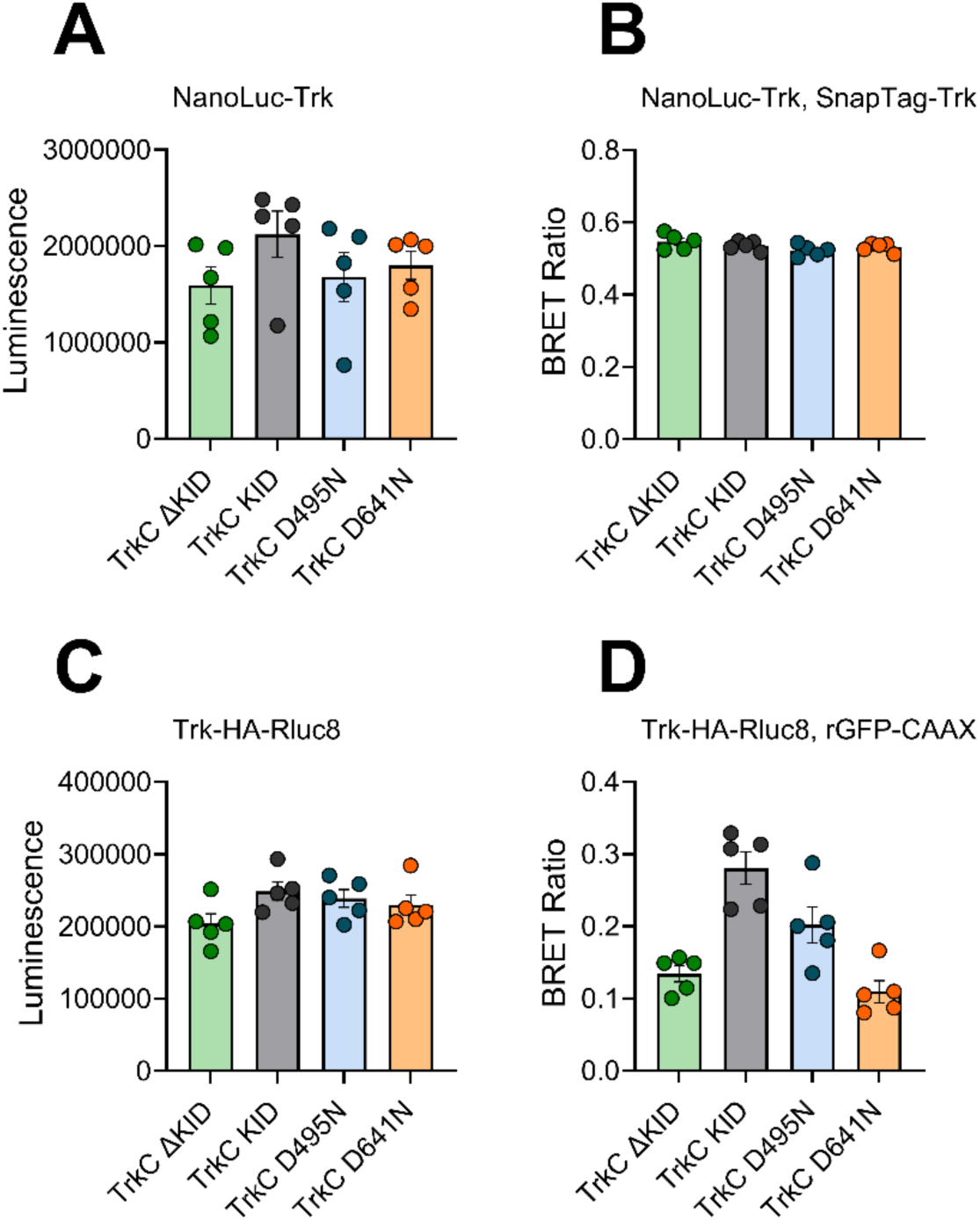
Luminescence emissions and pre-ligand addition BRET ratios for TrkC_KID_, TrkC_ΔKID_ or TrkC_ΔKID_ caspase cleavage mutants. (A) Receptor expression (Rluc8 luminescence emissions) and (B) BRET ratio between C-terminal tagged luminescent donor (Rluc8) and fluorescent acceptor (RGFP-CAAX) in the absence of ligand. (C) Receptor expression (NanoLuc luminescence emissions) and (D) BRET ratio between N-terminal tagged luminescent donor (NanoLuc) and fluorescent acceptor (SnapTag-AlexaFluor488). Mean ± SEM, from 4-5 independent experiments with triplicate wells.

